# Exploring the conformational space of a receptor for drug design: an ER*α* case study

**DOI:** 10.1101/2020.05.06.081000

**Authors:** Melanie Schneider, Jean-Luc Pons, Gilles Labesse

## Abstract

**Motivation:** Protein flexibility is challenging for both experimentalists and modellers, especially in the field of drug design. Estrogen Receptor alpha (ER*α*) is an extensively studied Nuclear Receptor (NR) and a well-known therapeutic target with an important role in development and physiology. It is also a frequent off-target in standard toxicity tests for endocrine disruption. Here, we aim to evaluate the degree to which the conformational space and macromolecular flexibility of this well-characterized drug target can be described. Our approach exploits hundreds of crystallographic structures by means of molecular dynamics simulations and of virtual screening.

**Results:** The analysis of hundreds of crystal structures confirms the presence of two main conformational states, known as ‘agonist’ and ‘antagonist’, that mainly differ by the orientation of the C-terminal helix H12 which serves to close the binding pocket. ER*α* also shows some loop flexibility, as well as variable side-chain orientations in its active site. We scrutinized the extent to which standard molecular dynamics simulations or crystallographic refinement as ensemble recapitulate most of the variability features seen by the array of available crystal structure. In parallel, we investigated on the kind and extent of flexibility that is required to achieve convincing docking for all high-affinity ER*α* ligands present in BindingDB. Using either only one conformation with a few side-chains set flexible, or static structure ensembles in parallel during docking led to good quality and similar pose predictions. These results suggest that the several hundreds of crystal structures already known can properly describe the whole conformational universe of ER*α*’s ligand binding domain. This opens the road for better drug design and affinity computation.

## 1 INTRODUCTION

In the physiological cellular environment, proteins are dynamic, which is often crucial for their function and activity. The conformational changes that a protein undergoes can range from atom vibrations or rotations of side-chains, to large shifts of secondary structures or domain reorientations, as well as changes in its multimeric organization. Theoretical models have been developed to simulate those conformational changes, but various limitations exist, such as the cost of lengthy simulations and approximations in the systems’ description [1]. Various biophysical characterizations of biological macromolecules have illustrated their intrinsic flexibility - at various scales in time (fs-s) and space (pm-nm). Nuclear Magnetic Resonance (NMR) spectroscopy was instrumental in highlighting various types of flexibility in atomic detail [2]. Unfortunately, this technique also suffers from several limitations (e.g. the limited size of the studied systems, or the need for multiple labelling) and yields usually lower resolution structures than X-ray diffraction. Crystal structures have been mostly viewed as static snapshots, shedding light on a small part of a macromolecule’s conformational landscape (e.g. end-point structures). However, for many therapeutic targets of interest the number of atomic structures has been growing, bringing a detailed view of their intrinsic variability, at least within the limit of crystal packing boundaries, which might represent a severe limitation in some cases. Meanwhile, refinement as structure ensembles demonstrated significant breathing even in crystal structures solved at temperatures as low as 100 K [3], although room temperature diffraction data seems to be required for more accurate modeling of the flexibility [4]. Interestingly, while some proteins show large structural rearrangements, it seems that most (potentially two-thirds [5]) experience only limited variations with low overall RMSD values (of less than 3 Å) between functionally relevant end-points. However, although it would have tremendous implications for ligand docking and affinity predictions, it is still unknown whether full coverage of the conformational space of folded proteins can be experimentally attained.

In parallel, structure-based virtual screening of small organic compounds has been mostly performed on single, rigid conformations of a target. However, docking accuracy falls off dramatically if an “average” or apo structure is used instead of an experimental crystal structure with a bound ligand [6]. Accordingly, docking new ligands requires to better describe structural variability. “Soft docking” uses reduced atomic volumes to allow small “overlaps” of the ligand and the receptor [7]. Obviously, this can introduce errors leading to false positives, and it also does not account for large conformational changes. Side-chain flexibility can be computed on-the-fly during the docking procedure using libraries of preferred conformers, as implemented in various docking programs, such as PLANTS [8]. Alternatively, the flexibility of a macromolecular structure can be pre-computed as an ensemble of static models. Such ensembles can be generated, for example, by molecular dynamics (MD) simulation [9, 10]. Some rearrangements can also be modeled using simpler calculations, such as normal mode analysis, at higher speed, but lower resolution [11]. MD simulations can also be performed after docking to address the flexibility of the complex, but this requires rather long MD simulations [12]. Parallel docking of a compound into multiple crystal structures may circumvent some of the above limitations by providing accurate structures of distinct conformations. This approach is evaluated here on a well-characterized therapeutic target with hundreds of crystal structures already published, as well as hundreds of ligands with high affinities.

The Estrogen Receptor alpha (ER*α*) is among the most studied Nuclear Receptors (NRs) [13]. Its Ligand Binding Domain (LBD) adopts the NR general fold of a three-layered *α*-helical sandwich comprising 12 helices (named H1 to H12) [14]. The ligand binding site is a mostly hydrophobic cavity buried in the core of the LBD. In the case of agonist-bound structures the ligand-binding cavity is sealed by the C-terminal helix H12. Antagonist-bound structures show a reorientation of the helix H12 to accommodate the usually larger antagonists. A third group of molecules, called Selective Estrogen Receptor Modulators (SERMs), shows mixed agonistic/antagonistic behavior depending on the tissue [15, 16, 17].

Numerous structures of the LBD of ER*α* have been solved in complex with various, but not all of its known ligands, and showed a more subtle variability beyond the large rearrangement of H12 [18, 19]. Nevertheless, it remains unclear whether all the accessible conformations of this LBD have been already observed. The wealth of well-characterized and high-affinity binders of this receptor were used to interrogate the conformation landscape in parallel to thorough structure comparison and focused numerical simulations of selected crystal structures.

## 2 APPROACH

In the present study the aim was to probe the complete conformational space of the nuclear receptor ER*α* based on freely available experimental data.The problem was approached from two complementary sides with different approaches: from the protein side by exploiting all crystal structures deposited in the PDB [20] in combination with free and restrained MD simulations (three techniques), and from the ligand side by making use of the known ligand space available via BindingDB [21].

First, all available crystallographic structures complexed with a ligand were analyzed as a structural ensemble to probe the flexibility of the receptor. Second, MD simulations were performed on three ER*α* complexes in agonist or antagonist conformations (PDBIDs: 2YJA, 2OUZ and 3UUC) to investigate the intrinsic dynamics and the ensemble generation capabilities. Third, re-refinement as structure-ensembles was also performed on the same three ER*α* crystal structures to interrogate the receptor flexibility in several crystal forms.

The resulting structural ensembles (MD simulations and X-ray ensemble refinement) of agonist 2YJA (complexed with estradiol) were submitted to Molecular Mechanics Poisson-Boltzmann Surface Area (MM-PBSA) affinity calculation in order to validate the generated ensembles from a protein-ligand interaction point of view.

Finally, two models of the receptor’s flexibility were challenged by docking all known high-affinity ligands of ER*α* (present in BindingDB).

The main questions was: Can we sufficiently cover the conformational space of our target receptor ER*α* to account for its interactions with ligands?

## 3 MATERIALS AND METHODS

### 3.1 Ensemble analysis of 440 ER*α* structures

All liganded ER*α* structures currently available in the PDB (461 protomers) were gathered using the @TOME server by submitting the canonical amino acid sequence of ER*α* (UniProt identifier: P03372) with a specified sequence identity threshold of 90%. 19 protomers (originating from 12 PDB entries) did not contain the C-terminal H12 and were therefore removed from the analysis dataset. Additionally, two outlier protomers (PDB-ID: 1A52) with an ambiguous electron density for H12, a rather low resolution (2.8 Å) and gold atoms, were also excluded for further analysis. The resulting structural dataset contained 440 ER*α* protomers.

All analysis and image generation was performed using the programming languages Python and R, in particular the R package ‘bio3d’ [22], and PyMOL (http://pymol.org).

### 3.2 Selected crystallographic structures

For MD simulation, ensemble refinement and local flexibility virtual screening (with PLANTS) representative structures of ER*α* were selected. As an agonist structure 2YJA was used, containing the natural ligand estradiol. Two antagonist structures were chosen, 2OUZ in complex with Lasofoxifene, a selective estrogen receptor modulator (SERM) and approved drug that is representative for antagonists in terms of size and shape, and 3UUC in complex with bisphenol C 2, an endocrine disruptor that represents the smallest pharmacophore structure as antagonist. These three crystal structures were also selected due to their good resolution (1.82 Å, 2.1 Å and 2.1 Å, respectively) and very good Diffraction Precision Index (0.14 Å 0.18 Å and 0.27 Å respectively) [23]. Structure 2YJA contains only one ER*α* monomer and no missing residues. Structure 2OUZ also contains only one ER*α* monomer, but several side-chains are missing. So, the complete structure was downloaded from PDB-REDO [24]. For 3UUC, the most complete monomer (chain D) contains gaps in four loop regions. Missing residues were added with an in-house script using Modeller [25], keeping the other residues fixed. Hydrogen atoms of ligands were added with OpenBabel at pH 7.

### 3.3 Molecular dynamics simulation

All simulations were carried out with Gromacs 2018 [10]. Ligand topologies were generated using the ACPYPE/ANTECHAMBER [26] program of AmberTools17 [27] with partial charges generated by the empirical charge model AM1-BCC. Ligand parameters were based on the General Amber Force Field (GAFF) while the Amber FF14SB force field was employed for the proteins. Each complex was solvated in a TIP3P water dodecahedral box, with periodic boundary conditions and a minimum distance of 1.0 nm from the surface of the complex to the edge of the box. Each system was neutralized by adding Na^+^ and Cl^−^ ions to physiological concentration of 150 mM. A completely free steepest descent energy minimization for 2000 steps was followed by a 100-ps NVT equilibration and a 100-ps NpT equilibration with Parrinello-Rahman pressure coupling at a reference temperature of 300 K and with ligand restraints of 1000 kJ/mol nm^2^ in x,y,z directions. Finally, 50 ns unrestrained production runs were performed with a 2 fs time-step in the NpT ensemble and snapshots were saved every 10 ps. For the simulations without ligands, the ligands were simply removed from the initial structures before starting the MD protocol. Analysis and plotting was performed with Gromacs tools, R and Python scripts.

### 3.4 Ensemble refinement of X-ray data

The Phenix tool `ensemble_refinement` models the experimental X-ray data by an ensemble of structures obtained by maximum-likelihood time-averaged restrained MD simulation [3]. Within the calculations, a large amount of sets of coordinates are sampled and the reported number of structures is reduced by selecting the minimal number of structures, equally distributed over the sampling time, that reproduces the *R*_free_ value of the whole trajectory within a 0.1% tolerance [3]. Diffraction data was downloaded from the PDB [20] and the ligand parameters were generated using Grade (http://grade.globalphasing.org). Two meta-parameters, *p_TLS_*, and *w_xray_* were varied (from 0.6 to 1.0 and from 2.5 to 10, respectively) following recommendations of the developers [3]. For the third parameter, the relaxation time *t_x_* (in ps) that changes the number of structures contributing to the target function, three values were chosen in relation with the diffraction resolution. In total, 45 separate simulations were performed for an exhaustive parameter test. The ensemble with the lowest *R*_free_ was selected for further analysis and was named, hereafter, an ensemble-refined structure.

### 3.5 MM-PBSA affinity calculations

For MM-PBSA affinity calculations, two types of conformation ensembles were used for estimating enthalpy energies by the single-trajectory MM-PBSA method implemented in g_mmpbsa [28]. Either classical 50-ns MD trajectories were reduced to 501 frames by extracting a frame every 100 ps. Alternatively, we used the ensemble-refined structures either directly used or after a short steepest-descent energy minimization. Calculations were performed based on a homogeneous medium with a dielectric constant of *ϵ_p_*=2 for the binding pocket and a value of *ϵ_s_*=80 for the solution, an ionic strength of 153.6 mM, an ionic radius of 0.95 Å for positive charged ions and 1.81 Å for negative charged ions, and a solvent probe radius of 1.4 Å.

### 3.6 Virtual screening

Known binders were extracted from BindinDB 2018 (http://bindingdb.org). This yielded 283 molecules with *Ki* affinity values. A last round of validation was performed using recently released BindingDB *Ki* dataset for ER*α*. Unfortunately, some compounds could not be modeled in 3D and could not be included in our docking study (e.g.: carborane or iron containing chemicals).

We attempted to dock all the known high-affinity ligands into the representative structures 2YJA and 3UUC using PLANTS [8]. To accommodate all the ligands into the latter, we allowed some side-chain flexibility in the binding site. Alternatively, we evaluated the use of multiple crystal structures in parallel to accommodate the same set of high-affinity ligands using the web-server @TOME2 as previously described [29]. Here, the similarity search was limited to structures retrieved using *Psi-Blast* and *HHSearch*, with a *90%* sequence identity threshold. For each ligand, 20 structures were selected according to ligand similarity (measured using the Tanimoto score as implemented in OpenBabel [30]). All the resulting screening campaigns are accessible here: http://atome4.cbs.cnrs.fr/Screening_ESRa

## 4 RESULTS AND DISCUSSION

### 4.1 Global flexibility analysis

#### 4.1.1 Structural variability of 440 crystal structures

To obtain a thorough picture of ER*α*’s intrinsic flexibility, 440 ER*α* protomers with 210 different co-crystallized ligands (originating from 232 PDB entries) were clustered and analyzed as a structural ensemble. Global flexibility was analyzed using clustering based on RMSD and Principal Component Analysis (PCA), ensemble Normal Mode Analysis (eNMA), RMSF analysis, and repeated for agonist and antagonist subsets.

PCA demonstrated an outstanding role of the first principal component PC1 which corresponds to H12 reorientation (Figure S3), with a huge proportion of the total variance (88.6%), compared to the second PC (1.65%). Similarly, eNMA and RMSF analysis pointed out, again, the high importance of the C-terminal H12 (Figures S5 and S6). Much smaller fluctuations were observed for the rest of the protein, as C*α* RMSF stays below 5.2 Å, with maxima at the N-terminus and the loops L11-12 (adjacent to H12) as well as L2-3 and L8-9 (numbered according to helix nomenclature).

Two different clustering methods (hierarchical and k-means) were employed using two different distance measures (RMSD and PC) (see Figure S1 and S2). All four clustering approaches partitioned the 440 structures into two main subsets, a large one with 358 protomers (agonists) and small one with 82 protomers (antagonists) (Figure S4). Within these two subsets, eNMA and RMSF analysis agreed and highlighted similar fluctuation peaks (Figures S5 and 1) for the same protein regions. Interestingly, agonist- and antagonist-bound structures showed flexibility in the same loops (L2-3, L8-9 and L11-12). However, agonist conformations showed highest fluctuations within loop L8-9 (and then L2-3), while antagonism involves higher fluctuations in loops L2-3 and L11-12.

**Figure 1.**
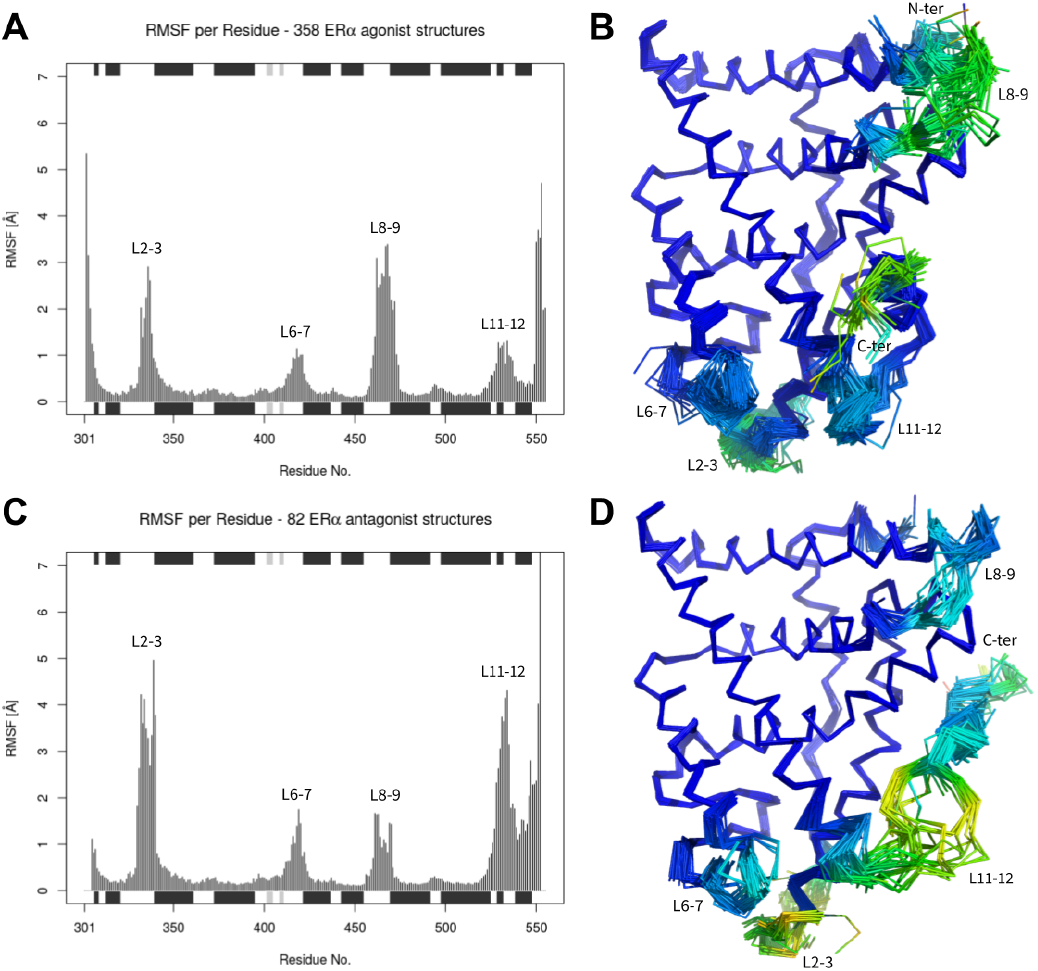
C*α* Root Mean Square Fluctuations (RMSF) of common-core superimposed ER*α* structures calculated on either the agonist subset (A and B) or the antagonist subset (C and D). Gap positions were included. The superimposed structures are colored by C*α* RMSF for the agonist subset (B) and the antagonist subset (D) (coloring scheme = rainbow, with a range of 0 to 7.39 Å).

Globally, our extensive structure comparison highlighted limited conformational rearrangements beyond that of H12 and some loops, although we cannot exclude crystallization artifacts in some cases. To address this issue, we characterized the flexibility of three representative crystal structures with conventional (unrestrained) MD simulations.

#### 4.1.2 Unrestrained MD simulations

The three systems investigated by 50 ns unrestrained MD simulations were the crystal structures bound to the natural agonist estradiol (PDB:2YJA) and to two antagonists, Lasofoxifene (PDB:2OUZ) and bisphenol C (PDB:3UUC). For these three systems, all five MD replicas showed stable and rather rigid behavior over 50 ns simulation time, as backbone RMSDs remained low along the trajectories with fluctuations of usually less than 2.5 Å (Figure S7). RMSF values averaged per residue, including backbone and side-chain atoms, showed baseline fluctuations in the 1 - 2 Å range (Figure 2 A and C, and Figure S6). The largest RMSF contributions came from H12 and flexible loop regions (compare Figure 2 B and D). While the mean RMSF per residue, averaged over all replica MD simulations, attained a maximum of 3.8 Å for the agonist 2YJA at residue Lys467 (located within loop L8-9), the antagonist 2OUZ attained a maximum of 8.6 Å at the C-terminus. When excluding H12 for 2OUZ, two mean RMSF peaks were visible with maxima of 4.5 Å at Phe337/Ser338 (within loop L2-3), and 4.6 Å at Lys531 (within loop L11-12).

**Figure 2.**
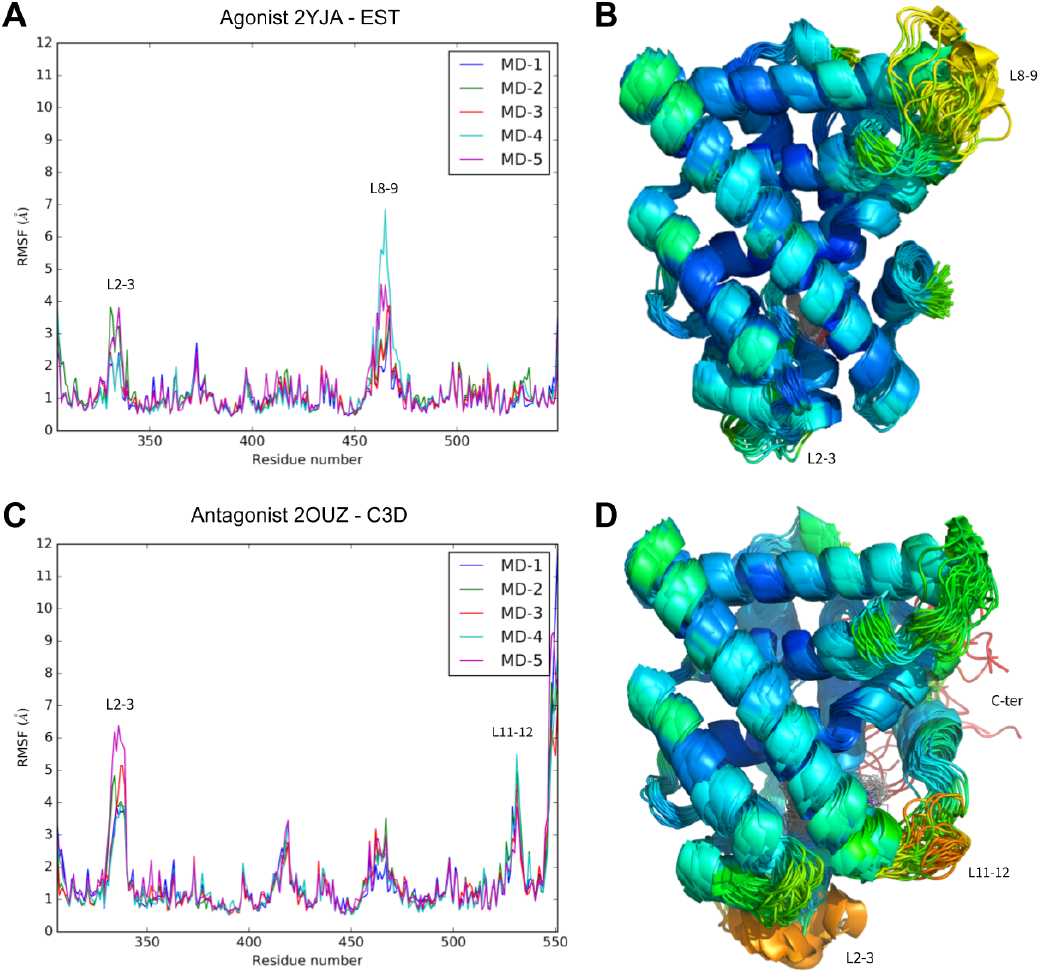
A) & C) RMSF averaged per residue of 5 50 ns MD simulations for agonist conformation 2YJA and antagonist conformation 2OUZ, respectively. B) & D) respective visualization of 55 frames extracted from MD-1 to MD-5 (one frame every 5 ns = 11 frames per MD) colored by RMSF average per residue averaged across all 5 MD simulations (coloring scheme = rainbow, with a range of 0 to 6 Å for B and D).

Repeating the simulations after removing the co-crystallized ligands did not reveal any significant changes in the global flexibility and behavior of the studied conformers on the timescale of the simulations (Figure S8). The overall agreement between the flexibility observed by unrestrained MD simulations and the variability among crystal structures prompted us to study the same selected crystal structures by ensemble refinement.

#### 4.1.3 Ensemble refinement: X-ray-restrained MD simulations

Ensemble refinement relies on very short MD simulations (a few hundreds of ps) restrained by X-ray diffraction data. It has revealed intrinsic flexibility in various crystal structures [3], but the biological relevance of such experimental structures is still debated. Our results suggested a distinct thermal fluctuation of the agonist form compared to the two antagonist forms studied here. Indeed, for 2YJA, the ensemble refined structure clearly showed an improved agreement with the experimental diffraction data (*R*_work_*/R*_free_ = 0.142/0.203) compared to the initial refinement as single model (*R*_work_*/R*_free_ = 0.198/0.234). On the contrary, for 2OUZ, ensemble refinement performed equally well (*R*_work_*/R*_free_ = 0.192/0.273) as a single-structure refinement (*R*_work_*/R*_free_ = 0.199/0.269). For 3UUC a more complex result was obtained, as the best ensemble showed only partial improvement (*R*_work_/*R*_free_ = 0.184/0.263) compared to the single model (*R*_work_*/R*_free_ = 0.214/0.255).

More refinement would be necessary to ascertain that crystal packing artifacts are not involved in the restrained conformation of the antagonist structures (although similar results were obtained with other antagonist-bound structures; see Table S1). Nevertheless, the flexibility observed in the refinement ensembles nicely matched that one deduced from crystal structure comparisons, as well as the one from unrestrained MD simulations. Interestingly, the ensemble-refinement explored even a slightly larger conformational space with larger RMSF values and larger variability in loop regions (compare Figures 3, S9, and S10).

**Figure 3.**
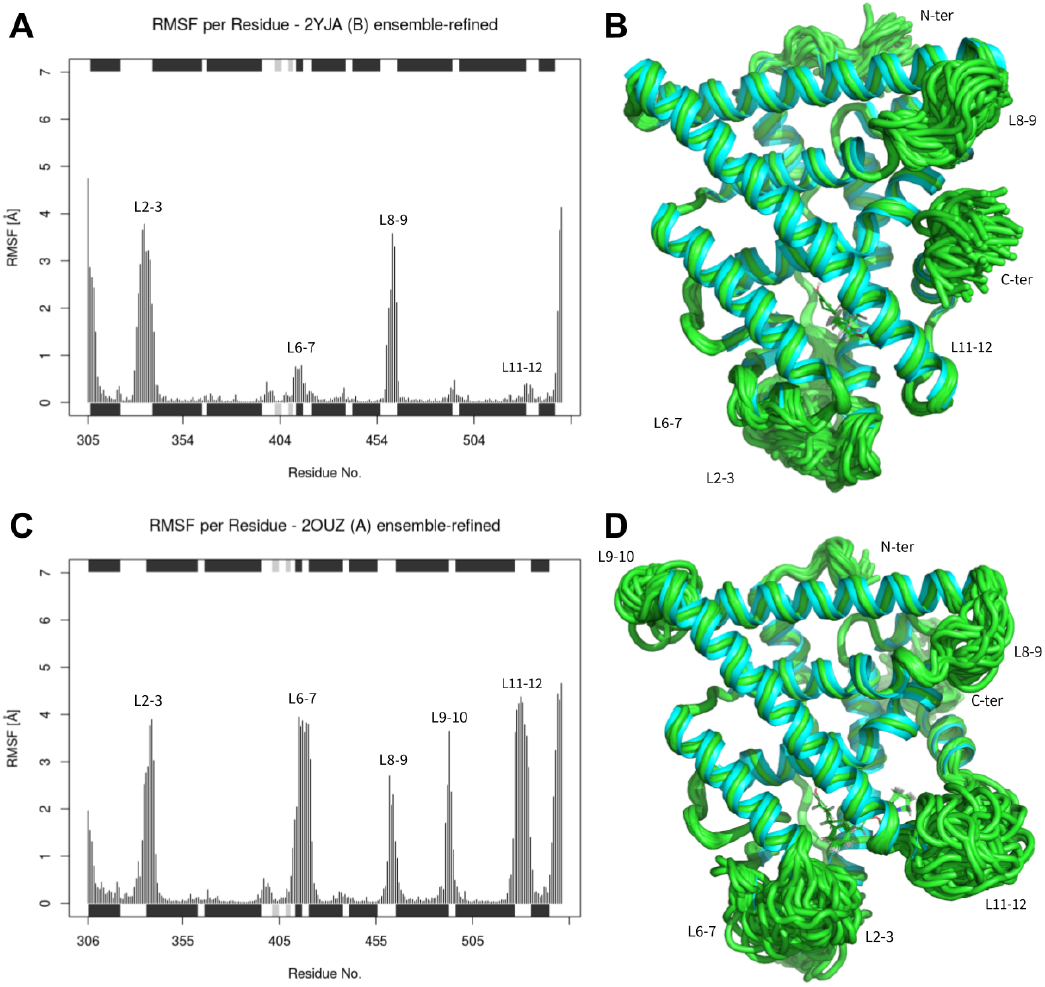
A) & C) RMSF averaged per residue of ensemble-refined agonist conformation 2YJA and antagonist conformation 2OUZ, respectively. B) & D) respective visualization of the single-refined (cyan) and the ensemble-refined (green) structures.

In this survey, the three methods (structure comparison, MD simulation, and ensemble refinement) revealed a similar level of global flexibility for ER*α* and showed very good overall agreement. This suggests that an ensemble of X-ray structures may reflect biologically relevant variability and intrinsic flexibility of this important medical target. This prompted us to focus into the binding site and its flexiblity.

### 4.2 Flexibility in the binding pocket

Here, the binding pocket of ER*α* is composed of 56 residues that were found at least once, at a 4 Å distance from a co-crystallized ligand. While, 20 residues very rarely participate in direct interactions, the remaining 36 residues can be attributed to three sub-pockets: a common one (with 18 residues very frequently (> 80%) in contact with a bound ligand), an antagonist-oriented one (11 residues) and an agonist-oriented one (7 residues) (see Figure 4A).

**Figure 4.**
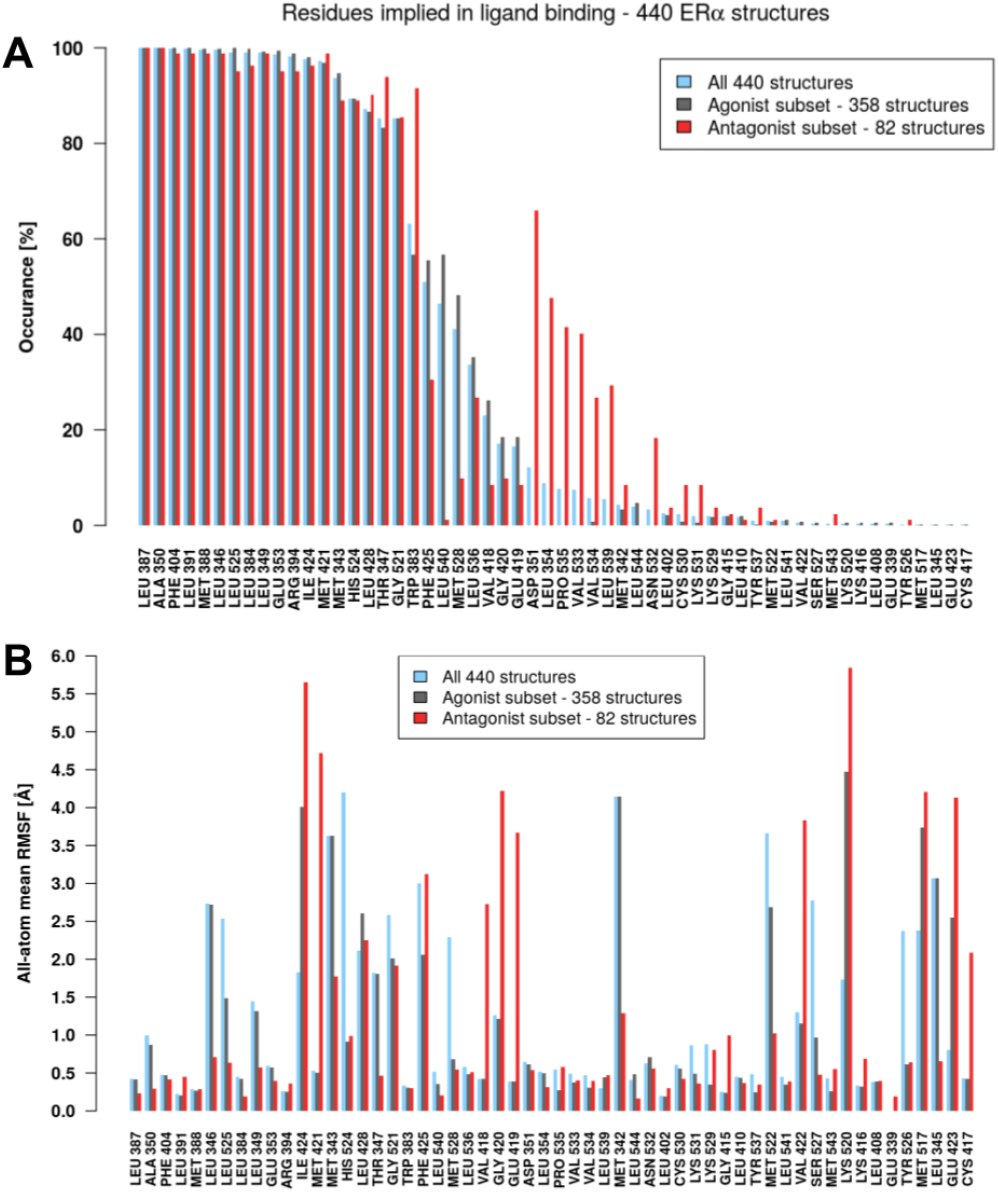
A) Frequency of involvement in ligand binding per residue (ordered from highest to lowest) for all 440 structures and for the conformer cluster subsets agonist and antagonist. B) All-atom mean RMSF per binding site residue, with residue ordering as in A.

Comparison of the 440 crystal structures showed small variations (< 0.5 Å) for half of the binding site residues, whereas rather large RMSF (of up to 5.5 Å) were detected for the remaining residues (e.g.: Ile424 and Lys520) (see Figure 4B). More precisely, significant fluctuations (> 2 Å) were observed for 23 residues, among which 8 were frequently ligand-contacting (e.g.: Ile424 and Gly521). This flexibility differed also depending on the agonist/antagonist nature of the ligand. Residues showing large agonist/antagonist differences included for example the frequently ligand-contacting Leu346, Leu525, Met421, and the segment Val418-Gly420.

In contrast, MD simulations showed a rather high background fluctuation for the whole binding pocket (no RMSF below 0.5 Å), with and without presence of a ligand during the simulation (see Figure 5A, S11, S12, and S13). It also highlighted increased flexibility for only 8 residues (> 2.5 Å), among which none was identified as frequent ligand contacting.

**Figure 5.**
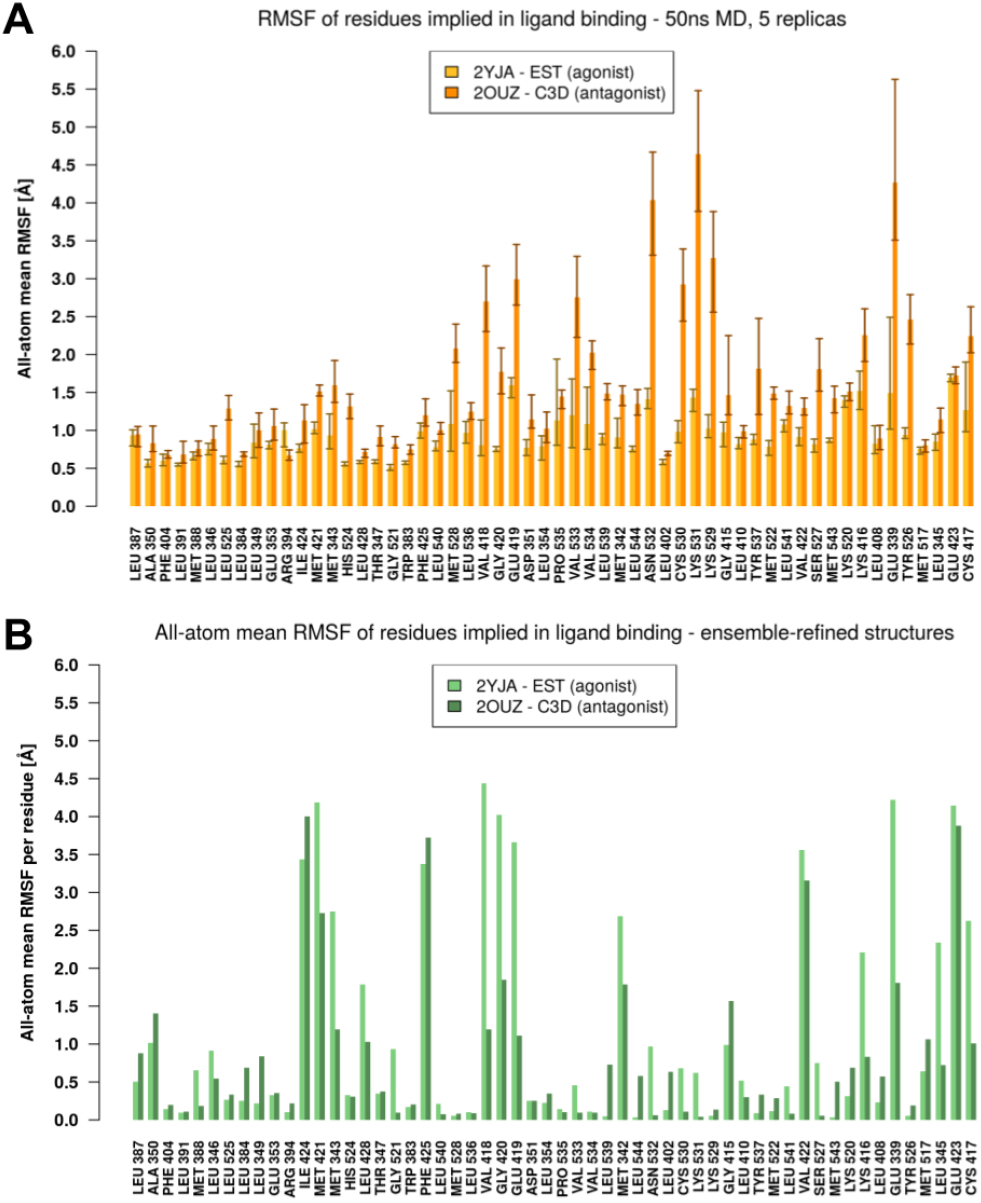
A) All-atom mean RMSF per binding site residue from 5×50 ns MD simulations, with residue ordering as in Figure 4. The height of the bars is the mean of the 5 replica simulations and the error bars indicate lowest and highest values within the 5 replicas. B) All-atom mean RMSF per binding site residue of the ensemble-refined crystal structures 2YJA and 2OUZ, with residue ordering as in Figure 4.

Ensemble refinement provided a picture more closely resembling that drawn from crystal structure comparison with respect to background fluctuations (1/3 below 0.5 Å) and flexibility peaks (compare Figure 4B, 5B and S14). This result suggests a more accurate representation of the protein flexibility within the binding site using very short simulation times (in the hundreds of ps instead of hundreds of ns) retrained by X-ray diffraction data.

### 4.3 MM-PBSA affinity calculations

To further evaluate the quality of the two distinct types of simulation (unrestrained and X-ray-restrained), we performed MM-PBSA computations to estimate precisely the enthalpic interactions between the ligand and the receptor for the example where flexibility was clearly observed (PDB:2YJA). Here, the classical computation on unrestrained MD snapshots showed a very favorable enthalpy term of about −270 kJ/mol of all the five replicas (Figure 6). Without energy minimization, the refinement ensembles had less favorable estimates of the enthalpic term (around −240 kJ/mol) and showed larger variations, including high-energy outliers that may correspond to protein conformations of high internal energy (Figure S15). Indeed, after standard energy minimization, the enthalpic term turned to be much more favorable (about −310 kJ/mol), with less variation between and within ensembles (Figure 6). We repeated the same MM-PBSA computation on the 20 complexes deduced from docking by @TOME (see below) for the estradiol ligand (Figure 6 and S16). Although it corresponds to a much lower number of conformations, the median enthalpy (about −260 kJ/mol) calculated on the minimized complexes fairly agreed with the one deduced from MD simulations.

**Figure 6.**
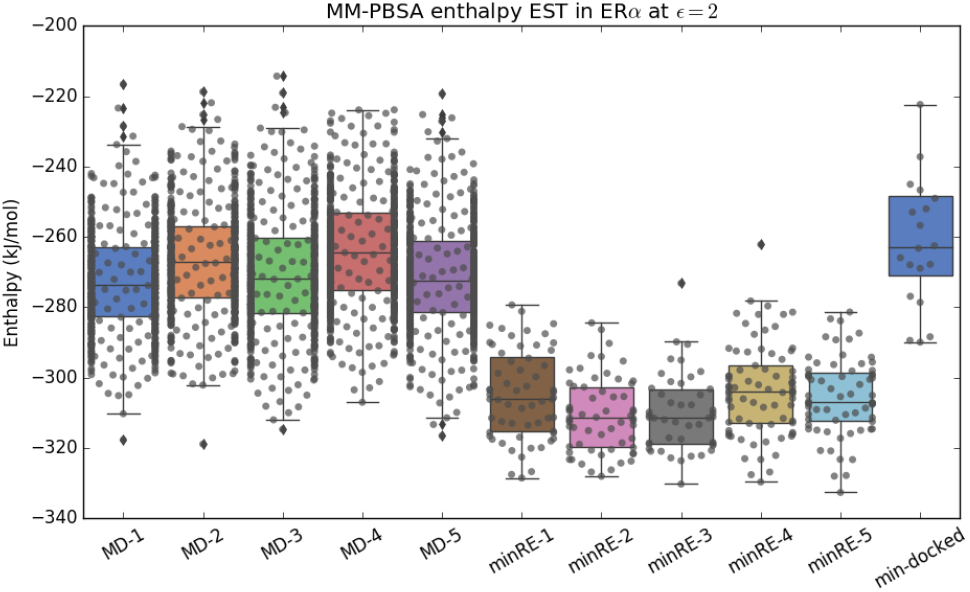
MM-PBSA calculations based on a) five replica 50 ns-MD trajectories (with 500 frames, one extracted every 100 ps) using PDB structure 2YJA that is complexed with estradiol (EST); b) the five best-*R*_free_ refinement ensembles (REs) (containing 30-55 structures each) using as well PDB structure 2YJA that is complexed with estradiol (EST); and c) docking results of estradiol (EST) into 20 different ER*α* PDB structures (selected by @TOME). The structures from b) and c) were minimized for 200 steps before calculation.

According to these results, minimized refinement ensembles or multiple docking complexes may provide a reasonable alternative to classical MD simulation as input to MM-PBSA calculations for affinity predictions. To evaluate further the current structural knowledge of this receptor, we aimed to produce convincing docking poses for all known ligands with sub-micromolar affinities.

### 4.4 Minimal required flexibility in the binding pocket

In order to probe which type of local flexibility is required for efficient virtual screening, systematic docking was performed on two selected crystal structures representative of the two main conformations (2YJA as agonist and 3UUC as antagonist) using PLANTS.

Small ligands, comprising most agonists and a few antagonists, could be docked successfully in a very similar and convincing mode into both selected protein conformations, the agonist (2YJA) and the antagonist (3UUC), as shown in Figure S17A. In contrast, ligands having bulkier extensions, and hence, mostly antagonist ligands, fit properly only into the binding pocket of the antagonist protein conformation 3UUC (compare Figure S17B and C). As one would expect, not all the ligands with high affinities could be docked successfully into 3UUC. From chemical similarities, shared by the accommodated ligands and those which were rejected, we identified some substitutions that could potentially clash with protein residues (by hypothetizing a similar binding mode). Therefore, two side-chains in the binding pocket were set as flexible, Met343 and Met421. Indeed, in this case, more ligands could be accommodated in the binding site with convincing orientations. Nevertheless, this was not sufficient for another class of large ligands, most of which were composed of a steroid core with a large extension connected to the D-ring via a rigid alkyne linker. A similar analysis highlighted two other side-chains in close proximity to the D-ring of agonists steroids, which were also set flexible (Met528 and His524). This resulted in successful docking of all the chemical series tested. The presented docking survey results are in agreement with the structural analysis described above (compare Figure 4B and S18). Indeed, in both cases a common set of side-chains was independently shown to be flexible for proper functioning of the receptor.

### 4.5 Virtual screening on static structure ensembles

As crystal structures harbor side-chains in favorable and variable conformations (see above), a second approach was used that attempted to automatically select a subset of optimal conformations for docking a given ligand. The target structure selection was based on ligand similarity (between docked and co-crystallized ligand), while the protein conformations were kept rigid during the docking (with PLANTS). We expected that this multi-structure-selection approach allowed to sample the available side-chain conformational space by using multiple experimentally determined X-ray structures. The same set of ligands were docked, and 70% of the automatically selected “best” poses (by the server @TOME) nicely matched our results from the manual screening. Most of the remaining ligands exhibited a convincing pose among the 19 other computed poses on @TOME, but were not automatically selected as “best” pose (http://atome4.cbs.cnrs.fr/Screening_ESRa/BindingDB2018). To further validate this approach, we also docked new ligands from the latest release of BindingDB. This included for example novel high-affinity compounds harboring a large adamantane moiety [31]. Their successful docking within the binding site of ER*α* confirmed the validity of our approach (http://atome4.cbs.cnrs.fr/Screening_ESRa/BindingDB2020). This suggested that the related ferrocene-containing compounds [31] can also be accommodated in known conformations of the receptor. Accordingly, we can readily dock all known binders into one or another crystal structures of the receptor already solved.

## 5 CONCLUSION

Comparison of the various available conformational ensembles of ER*α* (all crystal structures, classical MD simulations, and refinement ensembles) highlighted a very good overall qualitative agreement, as the same regions were found variable and flexible using the three approaches. Interestingly, the refinement ensembles appeared to recapitulate the side-chain flexibility better than classical MD simulations with respect to the variability found in the whole crystal structure set. Accordingly, they may represent an advantageous set for ligand docking and possibly MM-PBSA affinity calculations. For the first time, to our knowledge, an exhaustive virtual screening campaign has been performed to probe a receptor’s intrinsic flexibility. Remarkably, the array of automatically selected static ER*α* structures performed very well in the large-scale ligand docking campaign. Our overall results match a similar conclusion drawn from a previous study [32] on three other proteins, where selected crystal structures performed better than MD simulated conformations. The automated virtual screening on the @TOME-2 server represents a streamlined, labour-saving approach that benefits from the large set of known ER*α* structures and hence their intrinsic variability. Based on the rapidly increasing number of crystal structures solved today, we foresee that this overall approach will be suitable for numerous other important therapeutic targets.

## ACKNOWLEDGEMENTS

We thank Cathy Royer and Emmanuel Margeat for careful reading of the manuscript. We thank Muriel Gelin for providing us the script to automate ensemble refinement, as well as Matteo Paloni, Remy Bailly and Alessandro Barducci for their advice and computational resources.

## REFERENCES

[1] John L Klepeis, Kresten Lindorff-Larsen, Ron O Dror, and David E Shaw. Long-timescale molecular dynamics simulations of protein structure and function. Current Opinion in Structural Biology, 19(2):120–127, April 2009.

[2] Haydyn D. T. Mertens and Dmitri I. Svergun. Combining NMR and small angle X-ray scattering for the study of biomolecular structure and dynamics. Archives of Biochemistry and Biophysics, 628:33–41, August 2017.

[3] B. Tom Burnley, Pavel V. Afonine, Paul D. Adams, and Piet Gros. Modelling dynamics in protein crystal structures by ensemble refinement. eLife, 1:e00311, December 2012. 00092.

[4] James S. Fraser, Henry van den Bedem, Avi J. Samelson, P. Therese Lang, James M. Holton, Nathaniel Echols, and Tom Alber. Accessing protein conformational ensembles using room-temperature X-ray crystallography. Proceedings of the National Academy of Sciences, 108(39):16247–16252, September 2011. Publisher: National Academy of Sciences Section: Biological Sciences.

[5] Peter Cimermancic, Patrick Weinkam, T. Justin Rettenmaier, Leon Bichmann, Daniel A. Keedy, Rahel A. Woldeyes, Dina Schneidman-Duhovny, Omar N. Demerdash, Julie C. Mitchell, James A. Wells, James S. Fraser, and Andrej Sali. CryptoSite: Expanding the Druggable Proteome by Characterization and Prediction of Cryptic Binding Sites. Journal of Molecular Biology, 428(4):709–719, February 2016.

[6] Jon A. Erickson, Mehran Jalaie, Daniel H. Robertson, Richard A. Lewis, and Michal Vieth. Lessons in molecular recognition: the effects of ligand and protein flexibility on molecular docking accuracy. Journal of Medicinal Chemistry, 47(1):45–55, January 2004. 00293.

[7] F. Jiang and S. H. Kim. “Soft docking”: matching of molecular surface cubes. Journal of Molecular Biology, 219(1):79–102, May 1991. 00391.

[8] Oliver Korb, Thomas Sttzle, and Thomas E. Exner. Empirical Scoring Functions for Advanced ProteinLigand Docking with PLANTS. Journal of Chemical Information and Modeling, 49(1):84–96, January 2009. 00321.

[9] David A. Case, Thomas E. Cheatham, Tom Darden, Holger Gohlke, Ray Luo, Kenneth M. Merz, Alexey Onufriev, Carlos Simmerling, Bing Wang, and Robert J. Woods. The Amber biomolecular simulation programs. Journal of Computational Chemistry, 26(16):1668–1688, December 2005.

[10] Berk Hess, Carsten Kutzner, David van der Spoel, and Erik Lindahl. GROMACS 4: Algorithms for Highly Efficient, Load-Balanced, and Scalable Molecular Simulation. Journal of Chemical Theory and Computation, 4(3):435–447, March 2008. 07109.

[11] Olivier Sperandio, Liliane Mouawad, Eulalie Pinto, Bruno O. Villoutreix, David Perahia, and Maria A. Miteva. How to choose relevant multiple receptor conformations for virtual screening: a test case of Cdk2 and normal mode analysis. European biophysics journal: EBJ, 39(9):1365–1372, August 2010. 00055.

[12] A. Lavecchia and C. Di Giovanni. Virtual screening strategies in drug discovery: a critical review. Current Medicinal Chemistry, 20(23):2839–2860, 2013. 00063.

[13] B. Lawrence Riggs and Lynn C. Hartmann. Selective Estrogen-Receptor Modulators Mechanisms of Action and Application to Clinical Practice. New England Journal of Medicine, 348(7):618–629, February 2003.

[14] Vanessa Delfosse, Marina Grimaldi, Jean-Luc Pons, Abdelhay Boulahtouf, Albane le Maire, Vincent Cavailles, Gilles Labesse, William Bourguet, and Patrick Balaguer. Structural and mechanistic insights into bisphenols action provide guidelines for risk assessment and discovery of bisphenol A substitutes. Proceedings of the National Academy of Sciences of the United States of America, 109(37):14930–14935, September 2012. 00075.

[15] Hitisha K. Patel and Teeru Bihani. Selective estrogen receptor modulators (SERMs) and selective estrogen receptor degraders (SERDs) in cancer treatment. Pharmacology & Therapeutics, December 2017.

[16] Joan S. Lewis and V. Craig Jordan. Selective estrogen receptor modulators (SERMs): Mechanisms of anticarcinogenesis and drug resistance. Mutation Research/Fundamental and Molecular Mechanisms of Mutagenesis, 591(1):247–263, December 2005.

[17] Qin Feng and Bert W. OMalley. Nuclear Receptor Modulation - Role of Coregulators in Selective Estrogen Receptor Modulator (SERM) Actions. Steroids, 90:39–43, November 2014.

[18] Sathish Srinivasan, Jerome C. Nwachukwu, Alex A. Parent, Valerie Cavett, Jason Nowak, Travis S. Hughes, Douglas J. Kojetin, John A. Katzenellenbogen, and Kendall W. Nettles. Ligand-binding dynamics rewire cellular signaling via estrogen receptor-. Nature Chemical Biology, 9(5):326–332, May 2013.

[19] Jerome C Nwachukwu, Sathish Srinivasan, Yangfan Zheng, Song Wang, Jian Min, Chune Dong, Zongquan Liao, Jason Nowak, Nicholas J Wright, Ren Houtman, Kathryn E Carlson, Jatinder S Josan, Olivier Elemento, John A Katzenellenbogen, Hai-Bing Zhou, and Kendall W Nettles. Predictive features of ligand-specific signaling through the estrogen receptor. Molecular Systems Biology, 12(4):864, April 2016.

[20] Helen M. Berman, Tammy Battistuz, T. N. Bhat, Wolfgang F. Bluhm, Philip E. Bourne, Kyle Burkhardt, Zukang Feng, Gary L. Gilliland, Lisa Iype, Shri Jain, Phoebe Fagan, Jessica Marvin, David Padilla, Veerasamy Ravichandran, Bohdan Schneider, Narmada Thanki, Helge Weissig, John D. Westbrook, and Christine Zardecki. The Protein Data Bank. Acta Crystallographica. Section D, Biological Crystallography, 58(Pt 6 No 1):899–907, June 2002. 20928.

[21] Tiqing Liu, Yuhmei Lin, Xin Wen, Robert N. Jorissen, and Michael K. Gilson. BindingDB: a web-accessible database of experimentally determined proteinligand binding affinities. Nucleic Acids Research, 35(Database issue):D198–D201, January 2007. 00773.

[22] Barry J. Grant, Ana P. C. Rodrigues, Karim M. ElSawy, J. Andrew McCammon, and Leo S. D. Caves. Bio3d: an R package for the comparative analysis of protein structures. Bioinformatics (Oxford, England), 22(21):2695–2696, November 2006.

[23] Dinesh Kumar Kala Sekar, Gurusaran Manickam, S.N. Satheesh, P Radha, S Pavithra, K P. S. Thulaa Tharshan, John Helliwell, and K Sekar. Online-DPI: A web server to calculate the diffraction precision index for a protein structure. Journal of Applied Crystallography, 48:939–942, June 2015.

[24] Robbie P. Joosten, Fei Long, Garib N. Murshudov, and Anastassis Perrakis. The PDB redo server for macromolecular structure model optimization. IUCrJ, 1(Pt 4):213–220, May 2014. 00148.

[25] A. Sali and T. L. Blundell. Comparative protein modelling by satisfaction of spatial restraints. Journal of Molecular Biology, 234(3):779–815, December 1993. 09586.

[26] Junmei Wang, Wei Wang, Peter A. Kollman, and David A. Case. Automatic atom type and bond type perception in molecular mechanical calculations. Journal of Molecular Graphics & Modelling, 25(2):247–260, October 2006. 01693.

[27] D. A. Case, D. S. Cerutti, T. E. Cheatham, T. A. Darden, R. E. Duke, T. J. Giese, H. Gohlke, A. W. Goetz, D. Greene, N. Homeyer, S. Izadi, A. Kovalenko, T. S. Lee, S. LeGrand, P. Li, C. Lin, J. Liu, T. Luchko, R. Luo, D. Mermelstein, K. M. Merz, G. Monard, H. Nguyen, I. Omelyan, A. Onufriev, F. Pan, R. Qi, D. R. Roe, A. Roitberg, C. Sagui, C. L. Simmerling, W. M. Botello-Smith, J. Swails, R. C. Walker, J. Wang, R. M. Wolf, X. Wu, L. Xiao, D. M. York, and P. A. Kollman. Amber 2017, University of California, San Francisco, 2017. 00000.

[28] Rashmi Kumari, Rajendra Kumar, and Andrew Lynn. g mmpbsaA GROMACS Tool for High-Throughput MM-PBSA Calculations. Journal of Chemical Information and Modeling, 54(7):1951–1962, July 2014.

[29] Melanie Schneider, Jean-Luc Pons, William Bourguet, and Gilles Labesse. Towards accurate high-throughput ligand affinity prediction by exploiting structural ensembles, docking metrics and ligand similarity. Bioinformatics (Oxford, England), July 2019.

[30] Noel M. OLBoyle, Michael Banck, Craig A. James, Chris Morley, Tim Vandermeersch, and Geoffrey R. Hutchison. Open Babel: An open chemical toolbox. J Cheminf, 3:33, 2011. 00943.

[31] Shabnam Farzaneh and Afshin Zarghi. Estrogen Receptor Ligands: A Review (2013-2015). Scientia Pharmaceutica, 84(3):409–427, April 2016.

[32] David J. Osguthorpe, Woody Sherman, and Arnold T. Hagler. Exploring protein flexibility: incorporating structural ensembles from crystal structures and simulation into virtual screening protocols. The Journal of Physical Chemistry. B, 116(23):6952–6959, June 2012.

