## Supplementary materials for "Exploring the conformational space of a receptor for drug design: an ER*α* case study"

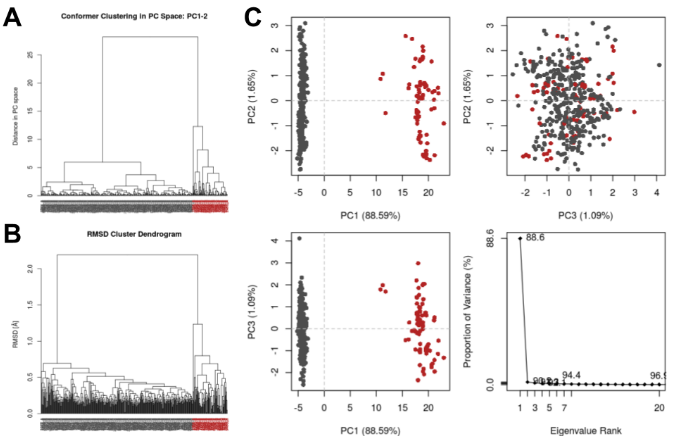


**Figure S1.** Hierarchical clustering of all 440 protomeric structures based on (A) distance in PC space and (B) RMSD. (C) Principal Component Analysis with plots for PC 1 to 3 and their respective proportion of variance. Coloring is based on RMSD cluster attribution.

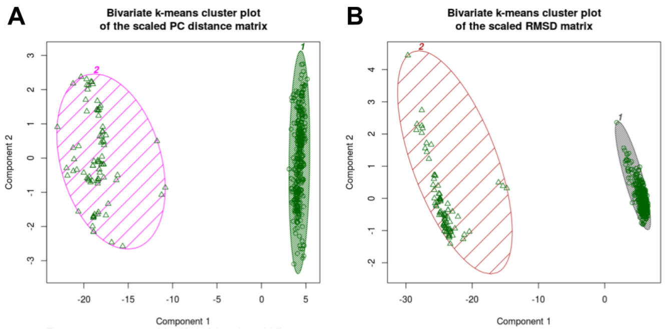


**Figure S2.** k-means clustering of all 440 protomeric structures based on (A) distance in PC space and (B) RMSD.

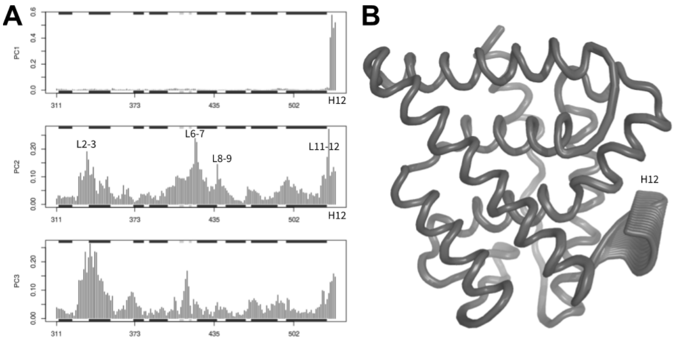


**Figure S3.** A) Principal Component (PC) residue contribution for the first three PCs and B) PC1 represented as trajectory.

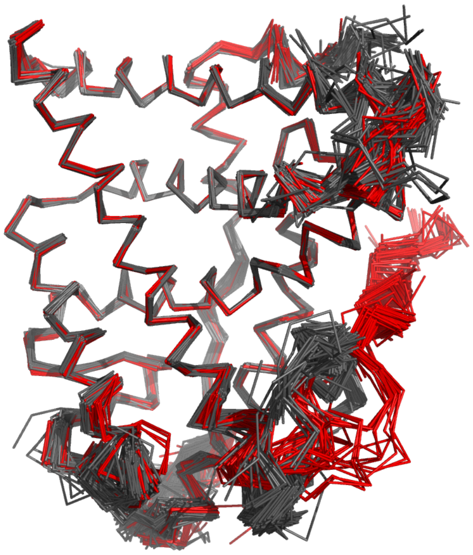

**Figure S4.** 440 superimposed ERα structures colored by conformer cluster - 358 agonist in grey and 82 antagonist in red.

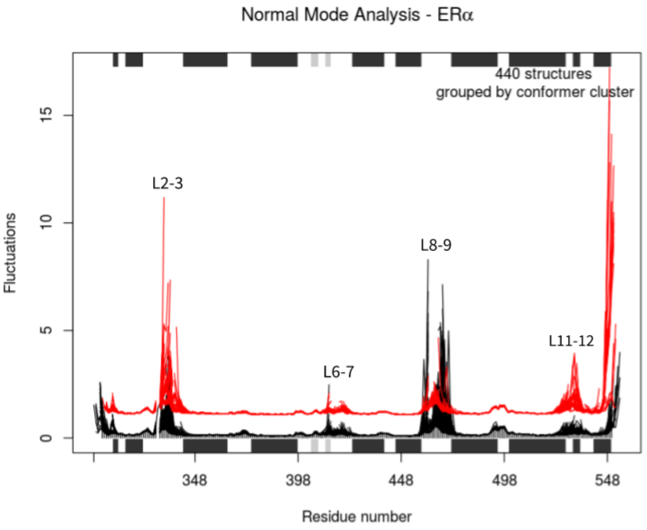


**Figure S5.** Ensemble Normal Mode Analysis (eNMA) with fluctuations per residue. The 440 protomeric structures were grouped by conformer cluster (agonist - black and antagonist - red) and spread for better comparison.

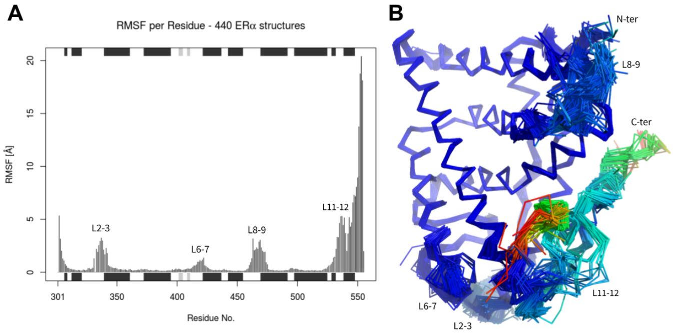


**Figure S6.** C$\alpha$ Root Mean Square Fluctuations (RMSF) calculated on all 440 protomeric ER$\alpha$ structures, superimposed on their common core C$\alpha$ atoms. A) C$\alpha$ RMSF plotted by residue with annotated secondary structures ($\alpha$ helix in dark grey, $\beta$ strand in light grey). Gap positions (residues not present in all structures) were excluded from the plot. B) Superimposed structures colored by C$\alpha$ RMSF including gap positions (coloring scheme = rainbow, with a range of 0 to 20.4 Å).

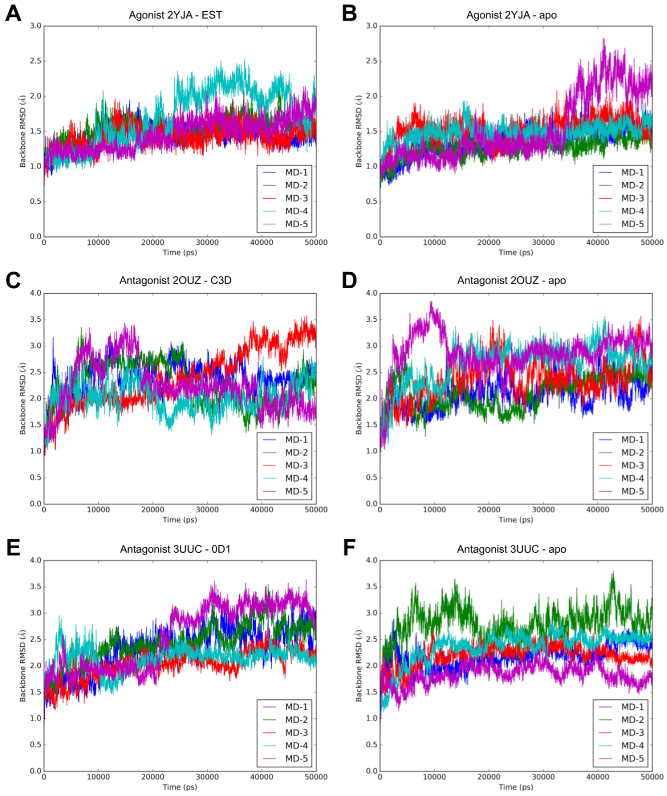


**Figure S7.** Backbone RMSD over time of 5×50 ns MD simulations for 2YJA (A), 2OUZ (C), and 3UUC (E), and of simulations without the respective ligands (as apo structures), (B), (D), and (F) respectively.

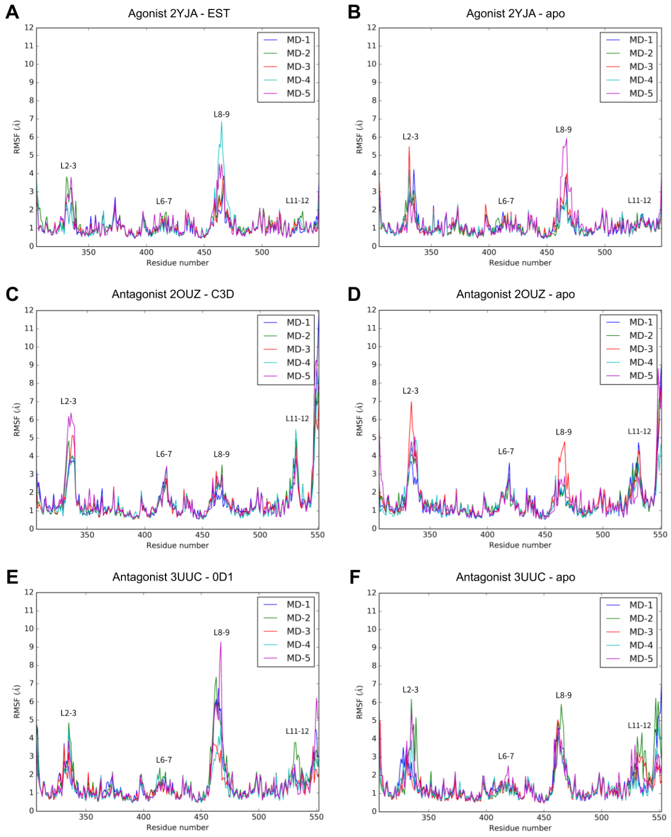


**Figure S8.** RMSF averaged per residue of 5x 50 ns MD simulations for agonist conformation 2YJA (A) and antagonist conformations 2OUZ (C) and 3UUC (E), and of simulations without the respective ligands (as apo structures), (B), (D), and (F) respectively.

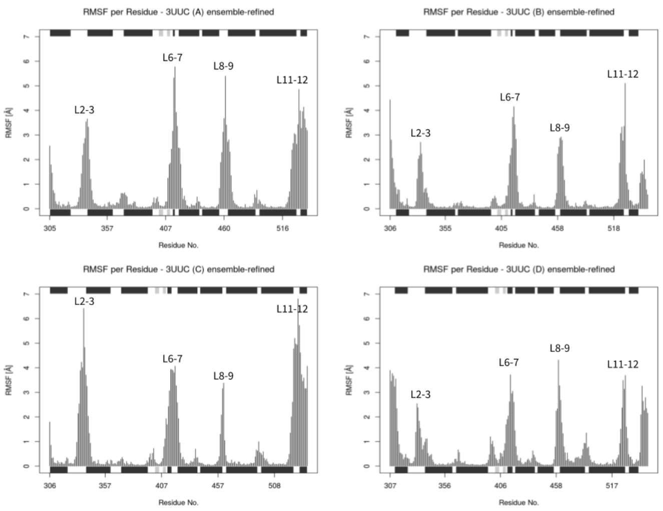


**Figure S9.** RMSF per residue for the four protomers (chain A-D) of the ensemble-refined crystal structure 3UUC.

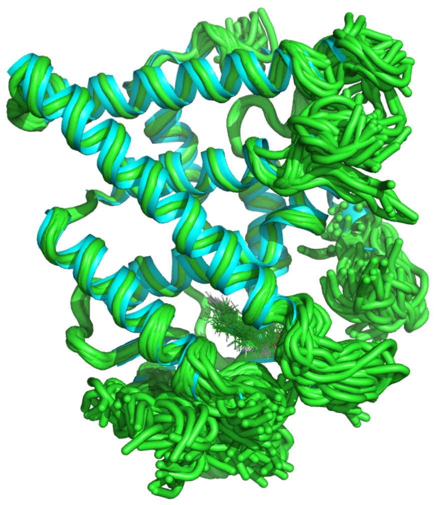

**Figure S10.** Protomer A of crystal structure 3UUC refined as single structure (cyan) and as ensemble (green).

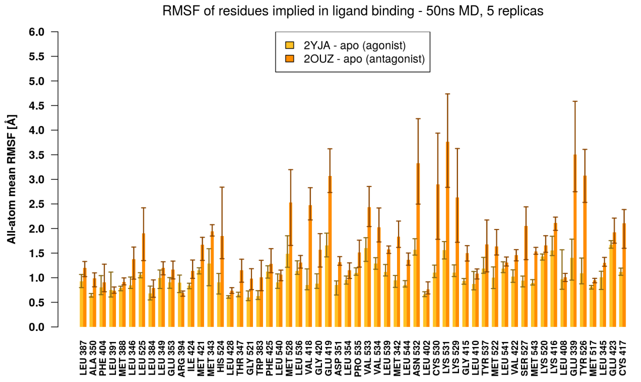


**Figure S11.** All-atom mean RMSF per binding site residue from 5×50 ns MD simulations, with residue ordering as in Figure 4. The height of the bars is the mean of the 5 replica simulations and the error bars indicate lowest and highest values of the 5 replicas.

**Table S1.** Selected ERα crystal structures, listed with PDB-ID, resolution and annotated ligand type. $R_{free}$ and $R_{work}$ are provided for the standard refinement as single structure and for the ensemble refinement.

| | | | single | | ensemble | |
|---|---|---|---|---|---|---|
| PDB-ID | Resolution | Ligand | $R_{work}$ | $R_{free}$ | $R_{work}$ | $R_{free}$ |
| 2YJA | 1.82 (Å) | agonist | 0.198 | 0.234 | 0.142 | 0.203 |
| 1GWQ | 2.45 (Å) | agonist | 0.193 | 0.261 | 0.158 | 0.232 |
| 5DID | 2.24 (Å) | agonist | 0.173 | 0.229 | 0.157 | 0.223 |
| 2OUZ | 2.00 (Å) | antagonist | 0.199 | 0.269 | 0.192 | 0.273 |
| 3UUC | 2.10 (Å) | antagonist | 0.214 | 0.255 | 0.184 | 0.263 |
| 5DWE | 1.92 (Å) | antagonist | 0.188 | 0.237 | 0.176 | 0.248 |
| 5UFW | 1.58 (Å) | antagonist | 0.169 | 0.201 | 0.167 | 0.218 |
| 4IVY | 1.59 (Å) | antagonist | 0.189 | 0.228 | 0.183 | 0.234 |
| 6VJD | 1.80 (Å) | antagonist | 0.188 | 0.224 | 0.173 | 0.232 |

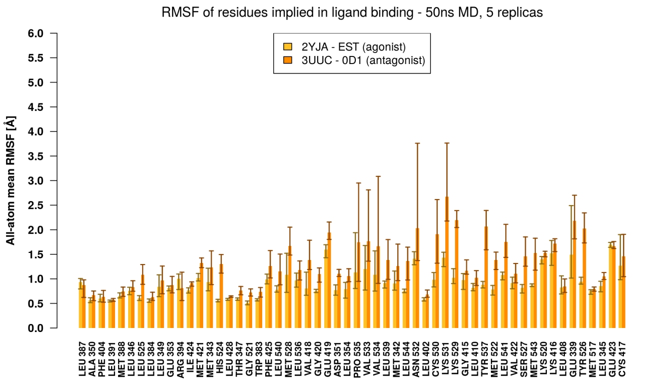


**Figure S12.** All-atom mean RMSF per binding site residue from 5×50 ns MD simulations, with residue ordering as in Figure 4. The height of the bars is the mean of the 5 replica simulations and the error bars indicate lowest and highest values of the 5 replicas.

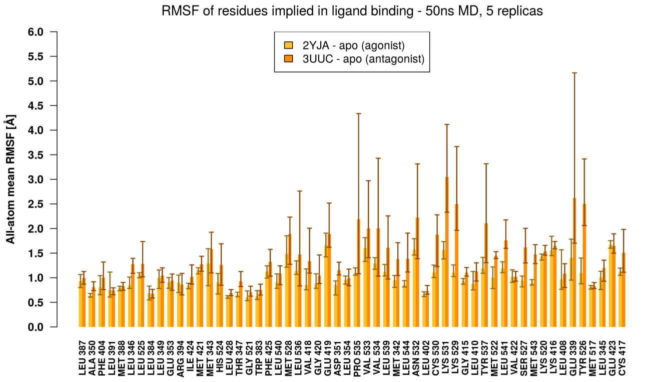


**Figure S13.** All-atom mean RMSF per binding site residue from 5×50 ns MD simulations, with residue ordering as in Figure 4. The height of the bars is the mean of the 5 replica simulations and the error bars indicate lowest and highest values of the 5 replicas.

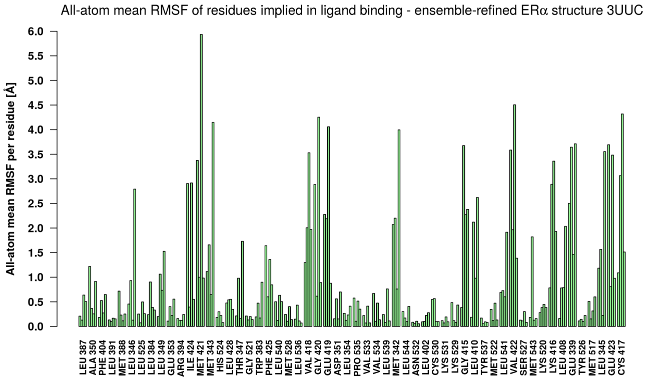


**Figure S14.** All-atom mean RMSF per binding site residue from the four protomers (chain A-D) of the ensemble-refined crystal structure 3UUC, with residue ordering as in Figure 4. The four protomer RMSF values were grouped per residue (four bars per residue for chain A-D).

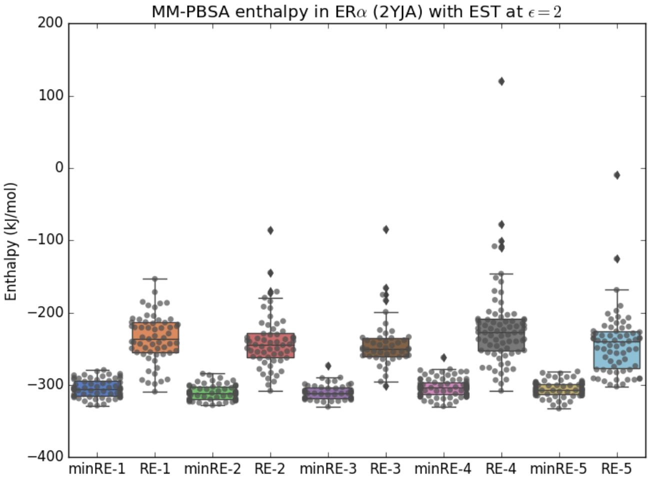


**Figure S15.** MM-PBSA calculations based on the five best-$R_{free}$ refinement ensembles (REs) (containing about 30-55 structures each) using PDB structure 2YJA that is complexed with estradiol (EST). The structures were either used directly for MM-PBSA calculation or minimized for 200 steps beforehand (indicated with 'min').

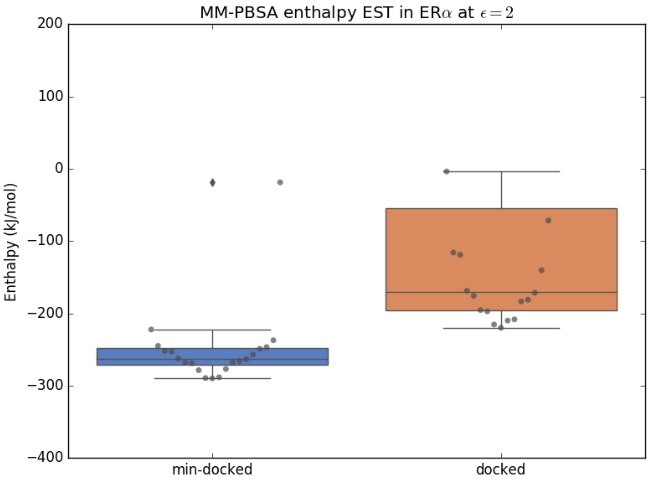


**Figure S16.** MM-PBSA calculations based on docking results of estradiol (EST) into 20 different ERα PDB structures (selected by @TOME). The structures were either used directly for MM-PBSA calculation or minimized for 200 steps beforehand (indicated with 'min').

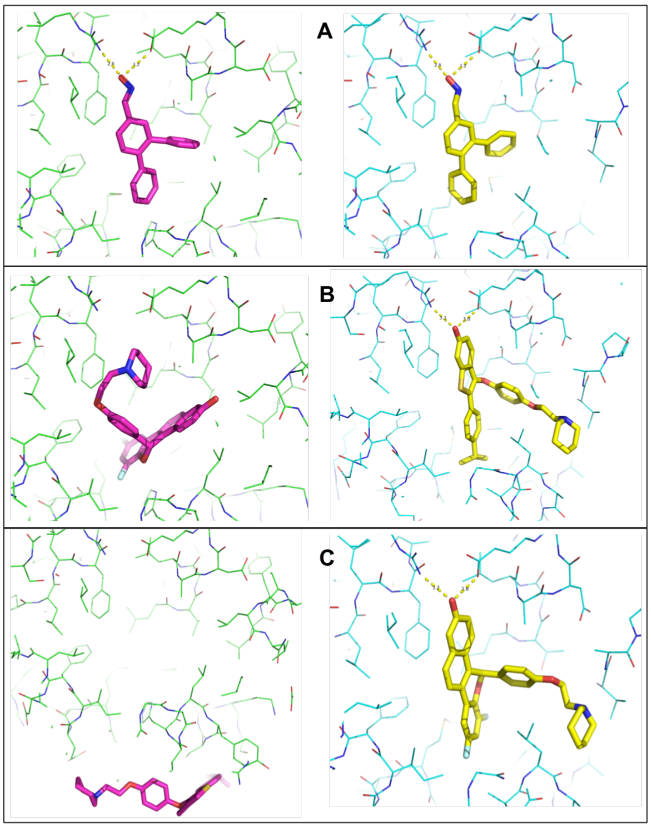


**Figure S17.** Exemplified docking results for three ERα ligands (A,B,C). The docked ligands are shown in purple (docked into 2YJA) and yellow sticks (docked into 3UUC), the protein structures are shown in green (2YJA) and cyan (3UUC) in line representation. Expected hydrogen bonds to Glu353 and Arg394 are depicted as yellow dotted lines. Ligand A shows convincing poses in both structures; whereat ligands B and C can only be docked successfully into 3UUC.

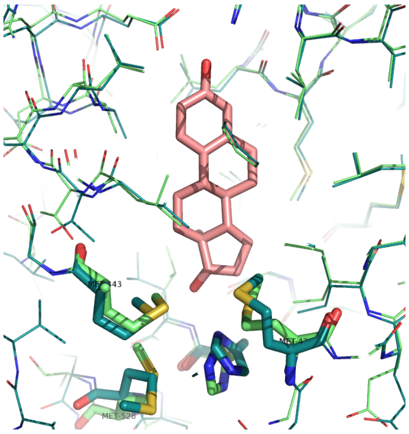

**Figure S18.** Comparison of side-chain conformations between structure 2YJA (in dark green) and 3UUC (in light green). The natural ligand estradiol is represented (in pink) for binding pocket localization. The selected four flexible side-chains (Met343, Met421, Met528 and His524), are located at the bottom of the binding pocket (highlighted in thicker stick representation).
